# Prior knowledge shapes neural routes to novel inference across events

**DOI:** 10.1101/2025.06.29.662181

**Authors:** Zhenghao Liu, Inês Bramão, Mikael Johansson

**Affiliations:** Department of Psychology, Lund University, Sweden

**Keywords:** memory integration, schema, context, MVPA, EEG

## Abstract

Memory allows us to go beyond direct experience by enabling inference across separate events. Yet how prior knowledge shapes the mechanisms supporting such inferences remains unclear. Here, we show that alignment between ongoing experience and existing knowledge determines how the brain constructs cross-event inferences. Behaviorally, inferences across schema-congruent events did not require accurate retrieval of the individual episodes, whereas inferences across schema-incongruent events depended on successful retrieval of those episodes. EEG-based multivariate pattern analysis further showed that schema-congruent inference was supported by schema reinstatement during encoding, consistent with the formation of integrated memory representations. In contrast, schema-incongruent inference relied on context-specific reinstatement at retrieval, reflecting the flexible recombination of distinct memories. Together, these findings demonstrate that prior knowledge dynamically shapes memory toward integration or separation, revealing how the brain constructs inferences to support flexible cognition.

## INTRODUCTION

Life events often share overlapping elements such as people, objects, and places, allowing memories to interact and enabling novel inferences across episodes (*1*). For example, seeing one person holding a book in a classroom and later observing someone else with the same book may lead to the inference that they study together. Although prior knowledge is known to support learning and memory (*2*), how it guides such cross-event inferences remains unclear. In particular, it remains unclear how the alignment of new experiences with prior knowledge—referred to as schema congruency—shapes the encoding of overlapping events and determines whether novel inferences arise from integrated representations or through the recombination of separate memories. Here, we address this question by combining electroencephalography (EEG) with multivariate pattern analysis (MVPA) to track neural representations across hierarchical levels (schema vs. context) and mnemonic stages (encoding vs. retrieval) during an inference task.

Memory-based inference relies on combining information across separate events and can arise through multiple mechanisms. One possibility is that familiar elements in new experiences trigger the reactivation of related past events during encoding, promoting the formation of integrated memory representations, combining old and new information, that support inference (*3–5*). Alternatively, to minimize interference, the brain may encode overlapping events as distinct representations that preserve episodic details, requiring inference to depend on the flexible retrieval and recombination of separate memory traces (*6–9*). Thus, whether memories are integrated or separated critically determines how inference is supported.

A key factor shaping memory formation is the extent to which new experiences align with prior knowledge. Schemas are abstract knowledge structures that capture regularities across experiences while filtering out perceptual detail, that provide a framework for organizing new information (*10*). For instance, although classrooms may vary in appearance, they share a common function as learning environments. Supported by the medial prefrontal cortex (*11*), schemas facilitate both encoding and retrieval (*12–15*). Importantly, their influence extends beyond congruent information. When new experiences violate expectations, prediction error signals can enhance memory, engaging the medial temporal lobe, particularly the hippocampus, to encode unique details and update existing knowledge (*16–20*). Together, these findings suggest that schema congruency exerts a non-linear influence on memory, with congruent and incongruent events engaging distinct neural processes (*2*).

Despite extensive evidence that prior knowledge shapes memory for individual events (*2*), it remains unclear how schema congruency influences inference across overlapping experiences. Specifically, it is unknown whether alignment with prior knowledge biases the brain toward integrating related events into a unified representation or maintaining them as separate memory traces. Addressing this question is essential for understanding how prior knowledge supports novel inferences.

To this end, we developed a paradigm in which participants encoded overlapping events while schema congruency was systematically manipulated (see **Figure 1A**). Participants first encoded AB events—picture-word pairs presented against a contextual background—with congruency determined by the semantic fit between the word and background (e.g., “desk” vs. “corn” in a classroom). A total of eight schemas were used (beach, classroom, city, kitchen, forest, airport, farm, and bathroom), each comprising eight contextual pictures (such as different beaches). Participants then encoded overlapping BC events, in which the same word was paired with a new picture against a neutral black background. Non-overlapping XY events were also encoded and served as a control. At test, participants performed inference on indirect AC associations, alongside memory tests for the associated schema and context, followed by memory tests for direct AB, BC and XY associations.

**Figure 1.**
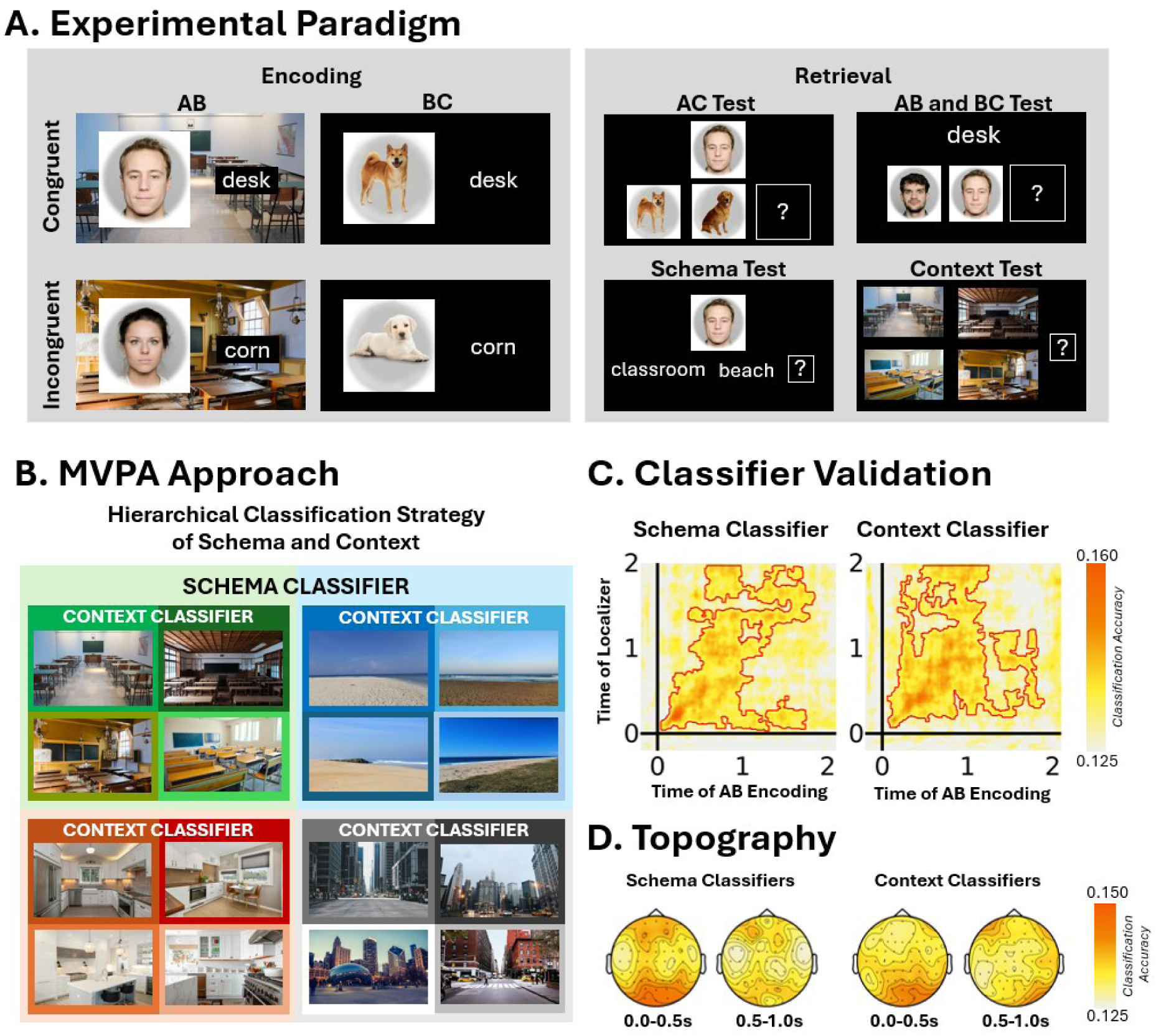
Overview of the paradigm and analytical approach. **(A) Experimental paradigm**. Participants encoded AB and BC events consisting of picture-word pairs presented on either a contextual (AB) or neutral (BC) backgrounds. Schema congruency in AB events was defined by the semantic fit between the word and the contextual background; for BC events, congruency was inherited from the corresponding AB pair. At retrieval, participants made inferences about indirect AC associations, and were tested on direct associations, and schema/context information. An “I don’t know” response was included to reduce noise from guessing. The face stimuli were selected from the Oslo Face Database (*29*), and individuals provided consent for their publication. Dogs and context images were similar to those used in the original experiment, but for illustrative purposes, we used license-free images from the Unsplash database (https://unsplash.com/license). **(B) Hierarchical classification approach.** EEG data from an independent localizer task were used to train hierarchical multivariate pattern classifiers to decode schema-level (eight semantic categories) and context-specific representations (individual exemplars within each schema). Context classification accuracy was averaged across exemplar classifiers. The figure displays four of the used schemas and four of their context exemplars. **(C) Time-generalized classification.** Classifiers trained on the localizer data were validated on AB encoding during periods when only the contextual background was presented. Classification accuracy above chance (12.5%) is indicated by red contours. Plots were smoothed with a 2D Gaussian kernel (50 ms width) solely for visualization purposes. **(D) Channel contribution maps.** Topographic maps show the contribution of individual EEG channels to schema and context classification.

To track neural representations, we trained hierarchical classifiers on EEG data from an independent localizer task to decode schema-level and context-specific information (see **Figure 1B**). These classifiers were validated during AB encoding and applied to BC encoding and AC retrieval. Previous research shows that reinstatement of an episodic event comprises both rapidly changing contextual details (*21–24*) and more stable, slowly drifting schematic representations (*25–28*). Building on this, we examined how schematic and contextual reinstatement vary as a function of schema congruency and contribute to inference.

We hypothesized that schema congruency determines which memory mechanism supports inference. When new information aligns with prior knowledge, shared schematic structure may promote the integration of overlapping events into a unified representation. In this case, reinstatement of prior events during BC encoding should support memory integration, reflected in schema-level reinstatement and predicting subsequent inference performance. In contrast, when new information violates prior knowledge, this mismatch may favor the encoding of events as distinct representations, preserving their unique details. Under these conditions, inference is expected to rely on the flexible retrieval and recombination of separate memory traces at test, reflected in context-specific reinstatement during retrieval.

## RESULTS

### Behavioral evidence for distinct mechanisms of inference

Participants (n = 39) performed an inference task requiring them to derive indirect AC associations from overlapping AB and BC events that were either schema-congruent or schema-incongruent. In addition to AC inference, we tested memory for schema and context information and for directly learned associations (AB, BC, XY) (**Figure 1A**).

We used a linear mixed model to test whether schema congruency predicted performance across all association types (see **Supplementary Note 1** for model equations). Participants were less accurate and slower when retrieving indirect AC associations compared with direct AB, BC, and XY associations (accuracy: *F*(3,38) = 105.668, *p* < .001, *η*ₚ² = 0.89; response times: *F*(3,36) = 99.096, *p* < .001, *η*ₚ² = 0.89), confirming that inference imposes additional cognitive demands. AB associations were retrieved more accurately and faster than BC and XY associations (all *p*s < .001), whereas BC and XY performance did not differ (all *p*s > .604). Schema congruency did not affect overall performance, with no main effect on accuracy (*F*(1,9824) = 0.164, *p* = .686) or response times (*F*(1,6121) = 0.153, *p* = .696). However, a significant interaction between congruency and association type was observed for accuracy (*F*(3,9824) = 3.509, *p* = .015, *η*ₚ² = 1.07 × 10^-3^), but not for response times (*F*(3,6107) = 1.328, *p* = .263). This interaction reflected reduced accuracy for schema-congruent compared with schema-incongruent BC associations (*t*(7368) = −2.392, *p* = .017, *d* = 0.043), indicating that schema congruency selectively impaired memory for BC associations without affecting overall inference performance (**Figure 2A**).

**Figure 2.**
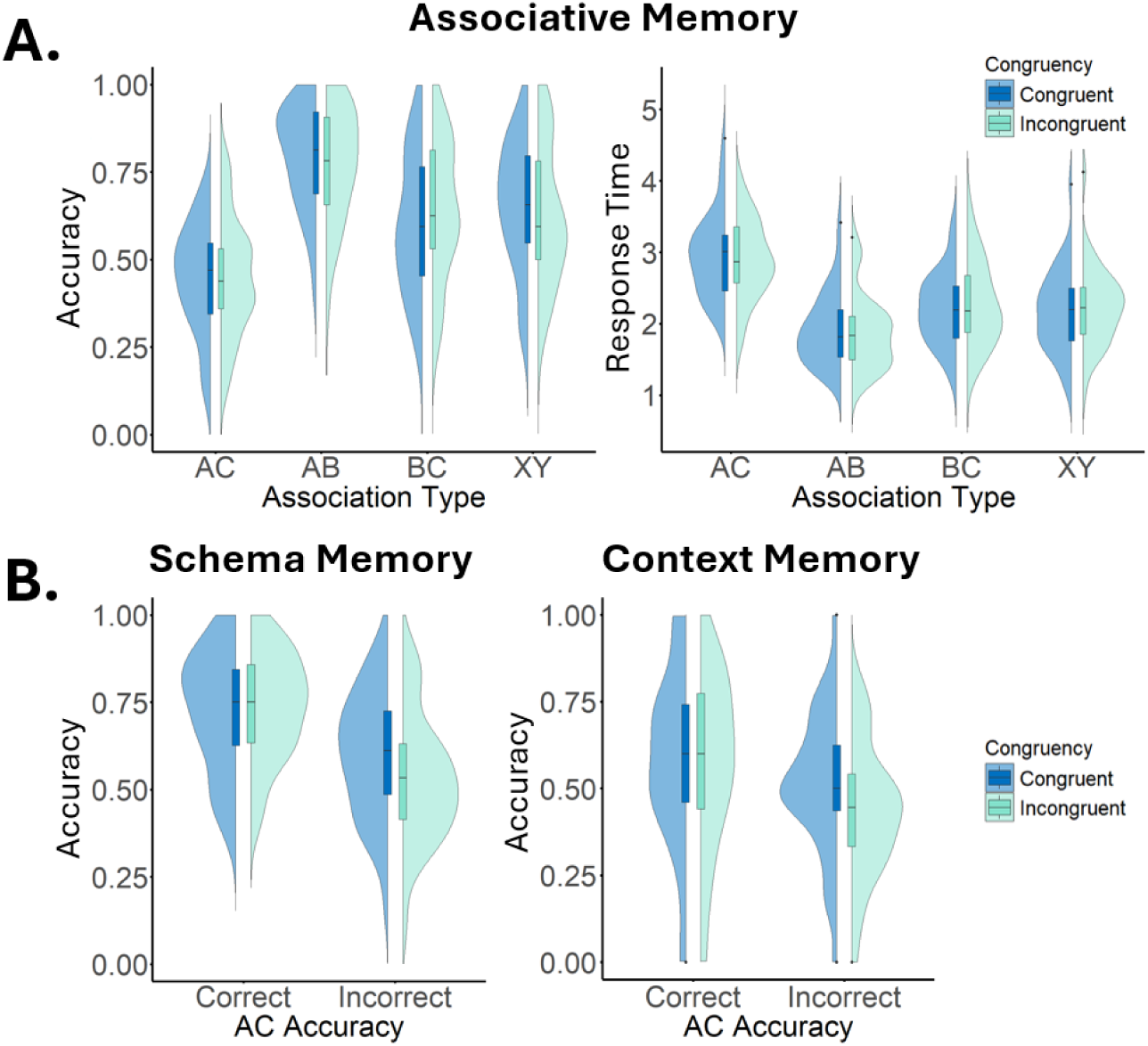
Behavioral performance for associative, schema, and context memory. **(A) Associative memory results.** Accuracy and response time for associative memory are shown as a function of Association Type (AC vs. AB vs. BC vs. XY) and Congruency (Congruent vs. Incongruent). **(B) Schema and context memory results.** Schema and context memory are shown as a function of AC inference accuracy (Correct vs. Incorrect) and Congruency (Congruent vs. Incongruent). Context memory was assessed only on trials with correct schema choice. Raincloud plots depict the distribution of individual data points, and the boxplots indicate the mean, and interquartile range.

To test whether inference relied on integrated or recombined memory representations, we examined whether AC performance was predicted by the joint retrieval of AB and BC associations (see **Supplementary Note 1** for model equations). If inference is supported by integrative encoding, AC performance should be independent of simultaneous AB and BC retrieval. In contrast, if inference relies on recombination, successful AC performance should depend on retrieving both AB and BC. Consistent with these predictions, AC inference for schema-congruent events was not predicted by the joint accuracy of AB and BC (*F*(1,1240) = 1.498, *p* = .221), suggesting that these events were integrated during encoding. In contrast, for schema-incongruent events, AC performance was predicted by the joint accuracy of AB and BC (*F*(1,1233) = 5.204, *p* = .023, *η*ₚ² = 1.21 × 10^-3^), indicating that inference relied on the retrieval and recombination of separate memory traces.

Together, these findings show that although schema congruency does not affect overall inference success, it fundamentally alters the computational mechanism supporting inference: schema-congruent events favor integrative encoding, whereas schema-incongruent events rely on retrieval-based recombination.

### Schema and context memory

We next examined schema and context memory as a function of schema congruency and inference success (**Figure 2B**). Schema memory was higher for schema-congruent than schema-incongruent events (*F*(1,2456) = 4.412, *p* = .036, *η*ₚ² = 1.79 × 10^-3^), whereas no effect of congruency was observed for context memory (*F*(1,1547) = 1.853, *p* = .174). Both schema and context memory were enhanced when AC inference was successful (schema: *F*(1,2495) = 86.783, *p* < .001, *η*ₚ² = 0.03; context: *F*(1,1571) = 21.117, *p* < .001, *η*ₚ² = 0.01), indicating that successful inference is associated with stronger memory representations. A significant interaction between congruency and inference success was observed for schema memory (*F*(1,2469) = 5.573, *p* = .018, *η*ₚ² = 2.25 × 10^-3^), but not for context memory (*F*(1,1552) = 2.345, *p* = .126). Post hoc comparisons revealed that when inference was incorrect, schema memory was higher for schema-congruent than schema-incongruent events (*t*(2466) = 3.328, *p* = .001, *d* = 0.082), whereas no congruency effect was observed when inference was correct (*t*(2467) = −0.184, *p* = .854; see **Figure 2**).

### Validation of hierarchical schema and context classifiers

To track schema-level and context-specific neural activity, we trained multivariate pattern classifiers on EEG data from an independent localizer task in which participants judged whether contextual pictures were indoor or outdoor (**Figure 1B**). Classifiers were trained to distinguish between schema-level and context-specific exemplars and were subsequently validated on EEG data from the main task.

Classifier performance was assessed during AB encoding in a time window in which only the contextual background was presented, allowing us to isolate schema and context representations independent of associative processing. Both schema and context classifiers showed reliable above-chance decoding, confirming that the classifiers captured meaningful neural representations (see **Figure 1C**).

Schema classifiers trained on localizer data between 0.1 and 1.9 s showed strong decoding along the diagonal, indicating temporally aligned representations. A similar pattern was observed for context classifiers, with diagonal decoding captured between 0.1 and 1.3 s. For both schema and context we observed additional off-diagonal generalization emerging between 0.8 and 2.0 s, reflecting broader temporal generalization (*30*) (**Figure 1C**). On averaged, classification accuracy exceeded chance level (12.5%) for both schema (mean accuracy ≈ 14%, BF₁₀ > 3.011) and context classifiers (mean accuracy ≈ 14%, BF₁₀ > 3.011), providing evidence for reliable decoding of both representational levels. These results indicate that both schema and context representations are reliably reinstated over time, with evidence for both temporally specific and temporally generalized neural patterns.

To characterize the spatial contributions to decoding, we examined channel contribution maps (*31*), see **Figure 1D**. Both schema and context classification show strong reliance on posterior channels, reflecting sensitivity to perceptual and context-specific information (*32*). Additionally, schema classification was associated with a frontal distribution of contributing electrodes, consistent with prior work implicating frontal regions in schematic processing (*10*, *33*).

Together, these results validate the use of hierarchical classifiers to dissociate schema-level and context-specific representations and establish a foundation for tracking their reinstatement during encoding and retrieval.

### Schema-congruent learning supports inference through memory integration

We next examined how schema and context representations are reinstated during encoding and retrieval for schema-congruent events. During BC encoding, the shared Word B was expected to trigger reinstatement of the previously encoded AB event, enabling the two events to be integrated into a unified memory representation that supports AC inference. We therefore tested whether schema and context reinstatement during encoding predicted subsequent AC inference performance (**Figure 3**).

**Figure 3.**
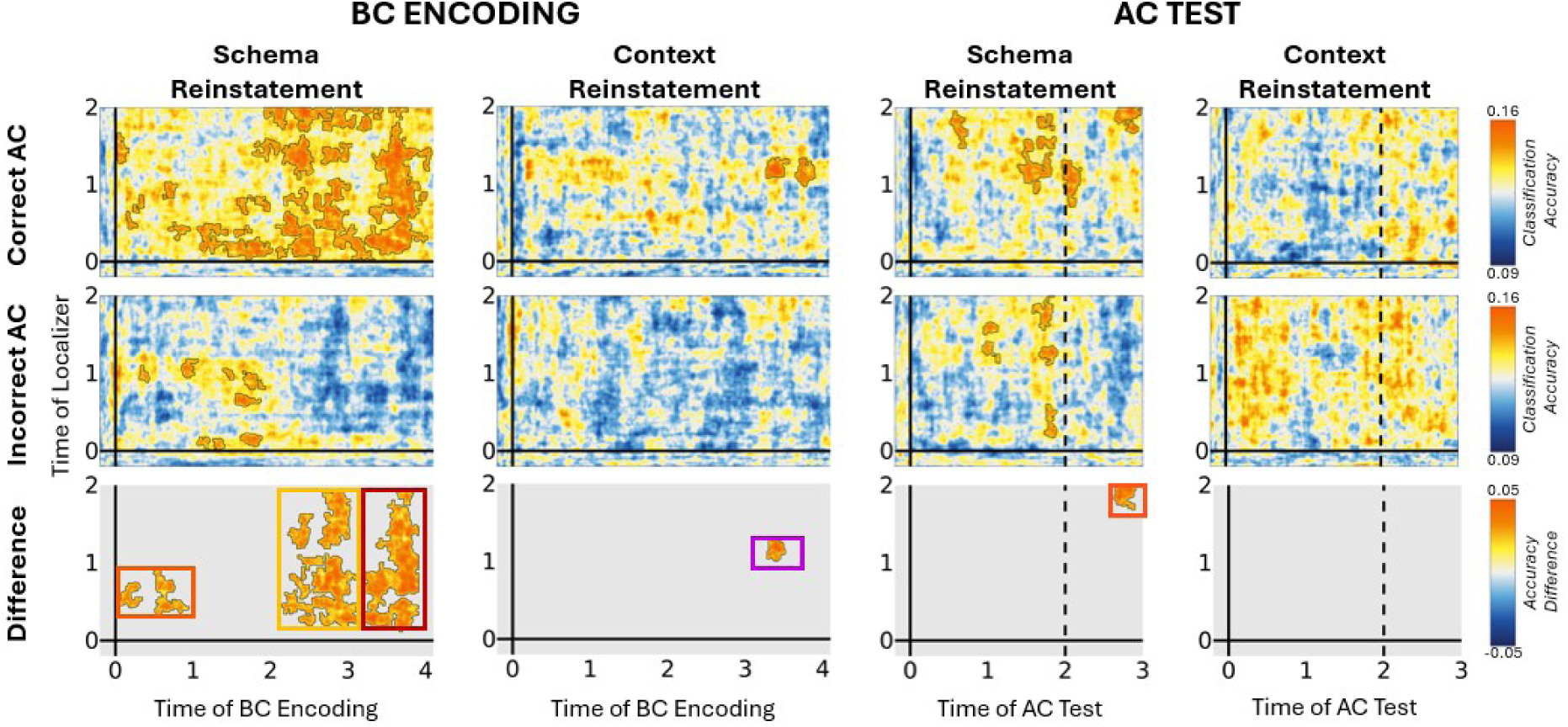
Schema and context reinstatement during BC encoding and AC retrieval for schema-congruent events. Trials with correct and incorrect AC inference are shown separately (top and middle rows) Blank contours indicate time points with classification accuracy significantly above chance. The bottom row shows the difference between correct and incorrect trials, highlighting reinstatement associated with successful inference. Time 0 corresponds to the onset of Word B and Picture C during BC encoding, and to cue onset during AC retrieval; the dashed line marks the onset of simultaneous presentation of cue, target, and distractor. Plots were smoothed with a 2D Gaussian kernel (50 ms) for visualization purposes.

Consistent with memory integration, schema reinstatement during BC encoding was associated with successful AC inference. Significant effects were observed in three time windows following the onset of Word B: 0.0-0.8 s (accuracy difference = 0.035, BF_10_ > 3.013), 2.5-3.1 s (accuracy difference = 0.037, BF_10_ > 3.012), and 3.2-4.0 s (accuracy difference = 0.038, BF_10_ > 3.012; **Figure 3**). These results indicate sustained reinstatement of schema-level information during encoding contributes to the formation of integrated representations linking AB and BC events. In addition to schema reinstatement, context reinstatement during BC encoding was also associated with successful inference, emerging later in the trial between 3.3 and 3.6 s (accuracy difference = 0.040, BF_10_ > 3.003; **Figure 3**). This late effect suggests that context-specific information can also contribute to encoding processes supporting inference. Our control analyses showed that schema reinstatement during BC encoding was not solely driven by the meaning of the on-screen word (see **Supplementary Note 3**). Compared with schema-congruent XY trials, which involve encoding without inference demands, schema-congruent BC trials elicited earlier and stronger schema engagement, indicating that this activity is driven by the need to make novel inferences.

At retrieval, we expected early schema reinstatement reflecting access to an integrated representation. Contrary to this prediction, schema reinstatement associated with successful inference emerged later in the trial, between 2.6 and 2.9 s, during the period in which the cue, target, and distractor were simultaneously presented on the screen (accuracy difference = 0.038, BF_10_ > 3.001; **Figure 3**). This delayed effect suggests that schema-level information contributes to later stages of inference processing rather than being immediately reinstated upon cue presentation.

To examine whether different forms of reinstatement reflect common or distinct processes, we analyzed their co-occurrence within trials with correct AC inference (**Figure 4A**). During encoding, early schema reinstatement predicted later reinstatement at 2.5-3.1 s (*β* = 0.233, BF_10_ = 7.855 × 10^5^), and both early and intermediate schema reinstatement predicted late reinstatement at 3.2-4.0 s (early: *β* = 0.133, BF_10_ = 314.956; intermediate: *β* = 0.458, BF_10_ = 1.787 × 10^26^), indicating a temporally sustained reinstatement process. In contrast, early schema reinstatement was negatively associated with context reinstatement during encoding (*β* = −0.092, BF_10_ = 7.738), suggesting a trade-off between schematic and context-specific processing. Similarly, schema reinstatement during encoding was negatively related to schema reinstatement at retrieval (*β* = −0.078, BF_10_ = 4.504), indicating that encoding- and retrieval-related processes may rely on partially distinct mechanisms. Finally, reinstatement strength was linked to memory performance. Late schema reinstatement during encoding (3.2-4.0 s) predicted subsequent schema memory (*β* = 0.054, BF_10_ = 4.446), whereas context reinstatement during encoding (3.3-3.6 s) predicted context memory (*β* = 0.104, BF_10_ = 38.076). These findings indicate that reinstatement at different representational levels contributes selectively to memory outcomes.

**Figure 4.**
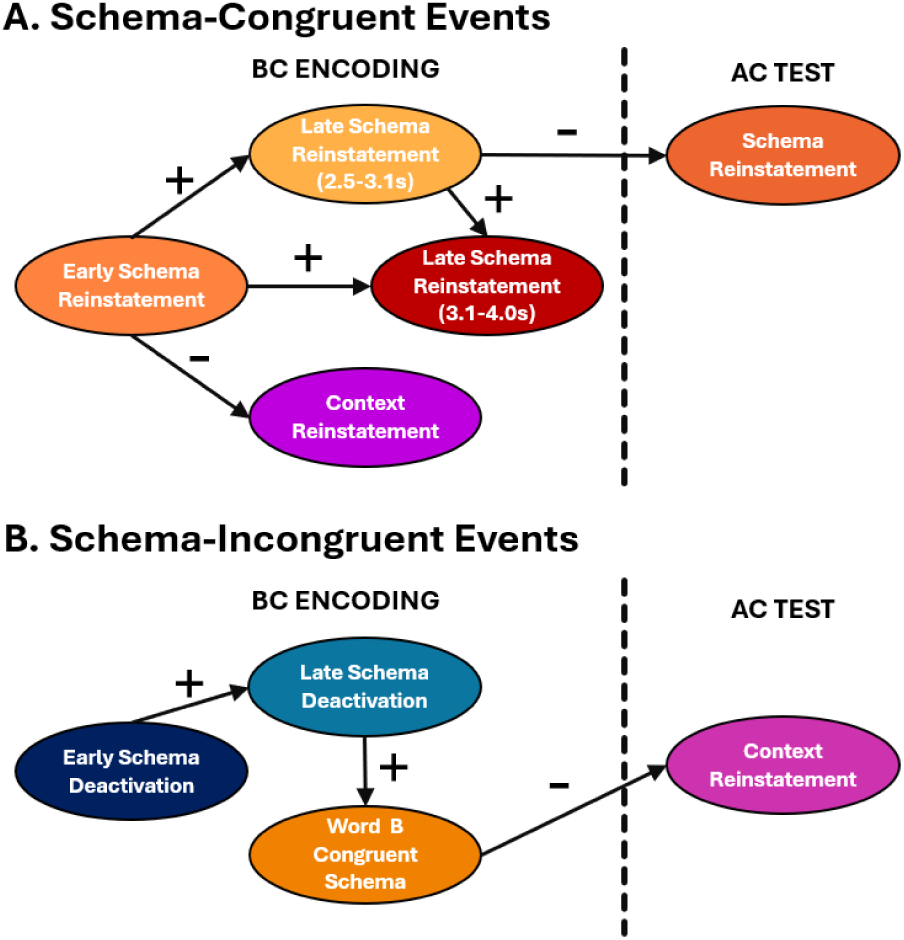
Co-occurrence of schema and context reinstatement associated with AC inference. Trial-level relationships between schema and context reinstatement are shown for **(A)** schema-congruent and **(B)** schema-incongruent events, measured during BC encoding and AC retrieval. Analyses were restricted to trials with correct AC. Only significant paths are shown.

Together, these results provide evidence that schema-congruent inference is supported by sustained schema reinstatement during encoding, consistent with integrative encoding mechanisms. When early schema reinstatement is weak or absent, later context reinstatement may also support successful inference, suggesting that multiple encoding processes can contribute. Notably, schema reinstatement at retrieval does not simply reflect the reactivation of an integrated representation but may instead support inference through distinct processes depending on the success of integrative encoding.

### Schema-incongruent learning shifts inference toward recombination of distinct memory traces

We next examined schema and context reinstatement during encoding and retrieval for schema-incongruent events. In contrast to schema-congruent conditions, the lack of alignment with prior knowledge was expected to reduce schema-based integration and instead favor the encoding of distinct memory representations. We therefore tested whether inference in this condition relied on context-specific reinstatement at retrieval, consistent with flexible recombination of separate memory traces (**Figure 5**).

**Figure 5.**
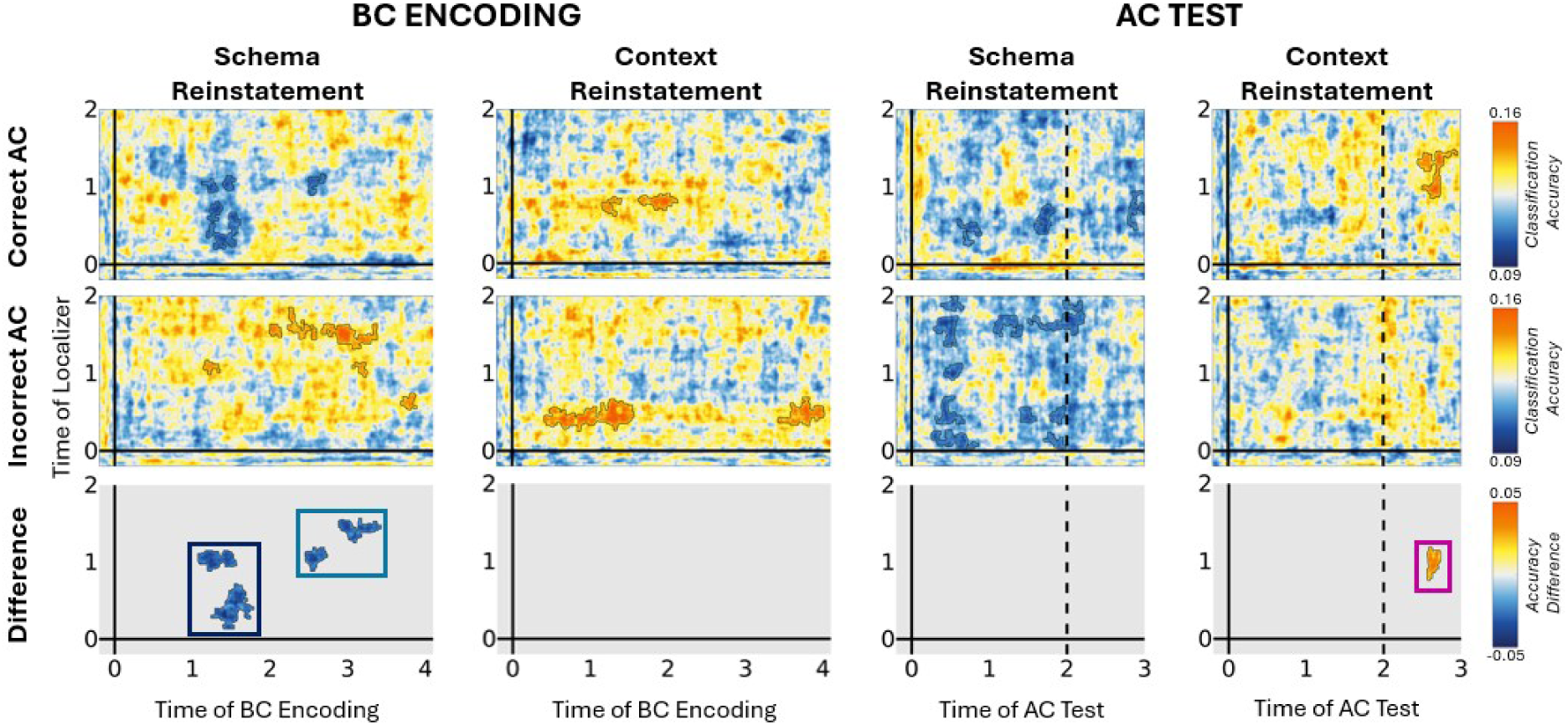
Schema and context reinstatement during BC encoding and AC test for schema-incongruent events. Trials with correct and incorrect AC inference are shown separately (top and middle rows). Blank contours indicate time points with classification accuracy significantly above chance. The bottom row shows the difference between correct and incorrect trials, highlighting reinstatement associated with successful inference. Time 0 corresponds to the onset of Word B and Picture C during BC encoding, and to cue onset during AC retrieval; the dashed line marks the onset of simultaneous presentation of cue, target, and distractor. Plots were smoothed with a 2D Gaussian kernel (50 ms) for visualization purposes.

During BC encoding, we did not observe schema reinstatement predictive of AC inference. Instead, schema classifier outputs revealed significant decreases in activation of the originally encoded (incongruent) schema in two time windows following the onset of Word B: 1.1-1.8 s (accuracy difference = −0.036, BF_10_ > 3.032) and 2.4-3.1 s (accuracy difference = −0.035, BF_10_ > 3.020; **Figure 5**). These results indicate suppression, rather than reinstatement, of the incongruent schema during encoding. Control analyses comparing BC and XY trials confirmed that this deactivation was not driven by perceptual or semantic mismatch between the word and schema (see **Supplementary Note 3**). Specifically, schema deactivation was observed during BC encoding but not during XY trials, indicating that it reflects processes related to inference demands rather than simple stimulus incongruency. To further characterize this effect, we examined whether schema deactivation was accompanied by the reinstatement of alternative schema representations (**Figure 6A**). Within time windows showing significant deactivation, classifier outputs revealed that the schema congruent with Word B was more frequently detected than baseline schemas during later stages of encoding (rate difference = 0.022, BF_10_ = 11.973). This suggests that suppression of the originally encoded schema is accompanied by the activation of a more relevant schema representation. Importantly, reinstatement of the Word B-congruent schema was greater for trials with correct AC inference than for incorrect trials (accuracy difference = 0.037, BF_10_ > 3.015; **Figure 6B**), indicating that schema-based processes can still contribute to successful inference under schema-incongruent conditions.

**Figure 6.**
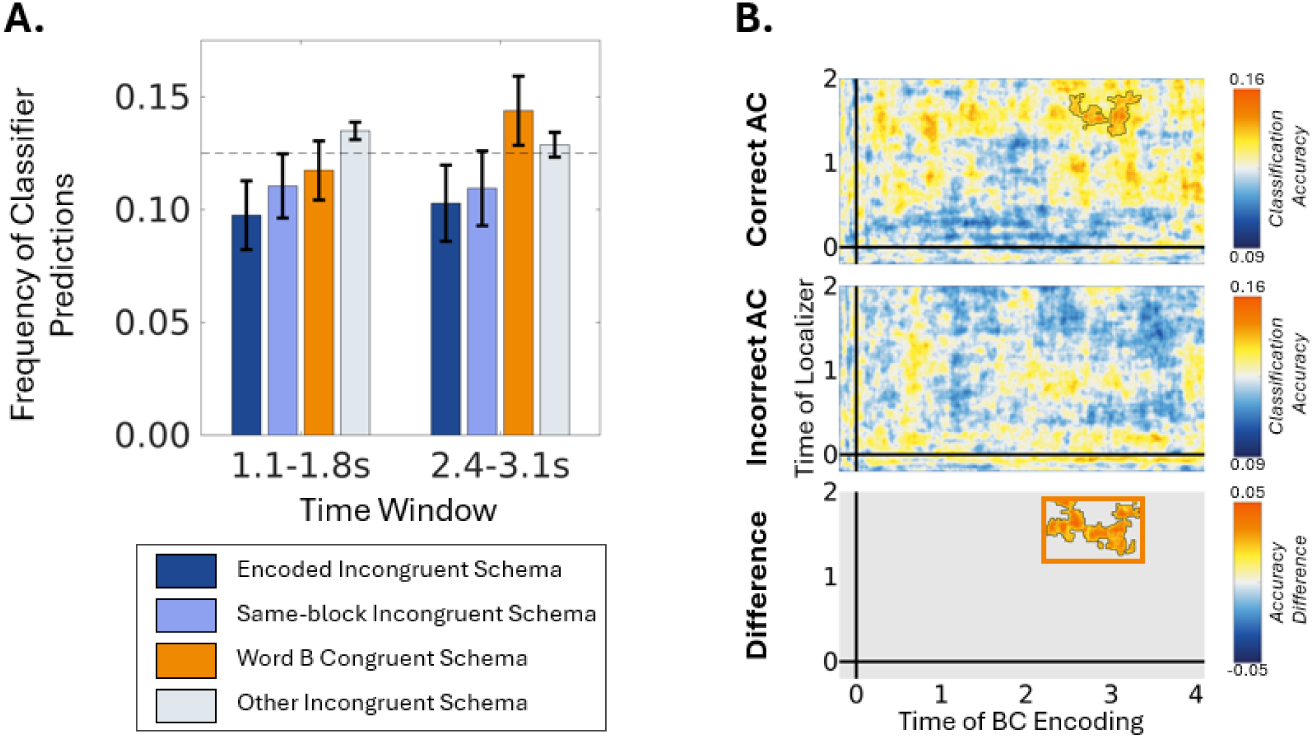
Reinstatement of alternative schema representations during schema-incongruent encoding. (A) Schema classifier outputs during BC encoding for the originally encoded (incongruent) schema, the same-block incongruent schema, the schema congruent with Word B, and all other schemas (baseline). (B) Reinstatement of the Word B–congruent schema during BC encoding as a function of AC inference performance, showing greater reinstatement for trials with correct inference. Plots were smoothed with a 2D Gaussian kernel (50 ms) for visualization purposes.

Consistent with a retrieval-based recombination account, context reinstatement during AC retrieval predicted inference performance. Significant effects were observed between 2.6 and 2.8 s (accuracy difference = 0.036, BF_10_ > 3.024), in a time window corresponding to the simultaneous presentation of cue, target, and distractor (**Figure 5**). This pattern reflects the reinstatement of context-specific information associated with the original AB event, which can be recombined with BC information to support inference. In contrast, no schema reinstatement predictive of AC inference was observed during retrieval (**Figure 5**), further indicating that inference in schema-incongruent conditions relies primarily on context-based processes rather than schema-level integration.

We next examined the co-occurrence of reinstatement effects within trials with correct AC inference (**Figure 4B**). Early schema deactivation during encoding predicted later deactivation (*β* = 0.392, BF_10_ = 2.694 × 10^19^), indicating a temporally sustained suppression process. In addition, late schema deactivation was associated with increased reinstatement of the Word B-congruent schema (*β* = −0.102, BF_10_ = 16.017), consistent with a shift toward more relevant schema representations. Interestingly, reinstatement of the Word B-congruent schema during encoding was negatively related to context reinstatement during AC retrieval (*β* = −0.065, BF_10_ = 3.136), suggesting that schema-based and context-based processes may provide complementary routes to successful inference rather than operating jointly within the same trials. Finally, we examined the relationship between reinstatement and memory performance. Context reinstatement during AC retrieval predicted context memory (*β* = 0.080, BF_10_ = 9.928), whereas no other reinstatement measures were associated with memory outcomes (BF_10_s < 1.724).

Together, consistent with our hypotheses, these results provide evidence that schema-incongruent inference is primarily supported by retrieval-based recombination of distinct, context-specific memory traces. Nonetheless, schema-level processes can still be engaged during encoding through the reinstatement of alternative, task-relevant schemas, and contribute to novel inferences.

## DISCUSSION

The ability to infer relationships across separate experiences is a fundamental feature of human memory, enabling flexible behavior beyond direct experience. Here, we show that prior knowledge systematically shapes the mechanisms supporting such inferences. Across behavioral and neural measures, schema congruency determined whether inference relied on integrative encoding or on the flexible recombination of distinct memory traces. These findings extend prior work on memory-based inference by demonstrating that the engagement of these mechanisms is not fixed, but dynamically modulated by the alignment between new experiences and existing knowledge.

Behaviorally, we observed that inference across schema-congruent events was not dependent on the joint retrieval of the underlying AB and BC associations, whereas inference across schema-incongruent events was dependent on their successful retrieval (**Figure 2**). This pattern aligns with prior work showing that memory-based inference can arise either through integrative encoding (*3–5*) or through retrieval-based recombination (*6–9*). Critically, our results go beyond this dichotomy by showing that prior knowledge influences which of these mechanisms is engaged. Schema-congruent experiences promote the formation of integrated memory representations that can be accessed directly at test, whereas schema-incongruent experiences preserve distinct episodic traces that must be retrieved and recombined to support inference.

Our neural findings provide converging evidence for this distinction and further clarify the underlying processes. A key contribution of the present study is the ability to track memory reinstatement across hierarchical representational levels using EEG-based multivariate pattern analysis (MVPA). By dissociating schematic and context-specific representations across both encoding and retrieval, this approach enabled us to examine how different levels of memory representation contribute to inference. Memory of naturalistic events is thought to be hierarchically organized, comprising rapidly shifting contextual details (*21–24*) and more slowly evolving and stable schematic knowledge (*25–28*). Building on this framework, our findings show that the relative dominance of these representational levels varies as a function of schema congruency. When new experiences align with prior knowledge, schematic representations are preferentially reinstated during encoding, supporting the formation of integrated memory representations that facilitate inference across events (*3–5*, *34*, *35*). In contrast, when experiences deviate from prior knowledge, contextual representations are emphasized, leading to the encoding of distinct memory traces that must be flexibly recombined at retrieval to support novel inference (*6–9*).

These findings are consistent with accounts proposing that schemas, supported by the medial prefrontal cortex, facilitate the organization and integration of new information (*10–15*). Prior work has shown that schema congruency enhances encoding and retrieval efficiency and supports the integration of related experiences into generalized representations (*11–15*). Extending this literature, our findings demonstrate that schematic knowledge not only facilitates memory but actively shapes the formation of representations that determine how subsequent inferences are made. Importantly, control analyses confirmed that the reinstatement of schematic knowledge reflects the demands of the inference task rather than mere responses to the semantic features of the presented Word B.

Additionally, schema reinstatement during BC encoding was associated with improved schema memory at retrieval, suggesting a reciprocal relationship: schemas facilitate encoding, while reactivating event content also strengthens memory for the schema itself. This finding is consistent with evidence that schema representations can be rapidly formed and remain stable over time, particularly when aligned with incoming information (*13*, *36*).

Successful inference for schema-congruent events was also associated with schema reinstatement during the AC retrieval. While such schema reinstatement could indicate retrieval of an integrated ABC representation, we observed a negative correlation between schema reinstatement during retrieval and BC encoding. This suggests that when an integrated representation is not formed during encoding, schema reinstatement at test may support inference by facilitating the flexible recombination of separate episodes. Conversely, when an integrated representation is available, schema reactivation at retrieval may not be necessary.

In contrast, when events were schema-incongruent, we observed schema suppression during BC encoding rather than reinstatement. This finding is consistent with theories proposing that violations of prior knowledge engage prediction error mechanisms that enhance the encoding of novel, context-specific information (*16–19*). Such processes have been linked to hippocampal encoding of distinct episodic representations and updating of existing knowledge structures (*16*, *17*). Our findings extend this work by showing that schema incongruency biases the system toward separation rather than integration of overlapping experiences, thereby shaping how subsequent inference must be constructed.

Importantly, inference in schema-incongruent conditions was supported by context reinstatement during retrieval, indicating reliance on the flexible recombination of distinct episodic memories. This is consistent with prior work demonstrating that inference can be constructed at test through the retrieval and recombination of separate memory traces (*6–9*). By linking this process to schema incongruency, our results suggest that the absence of an organizing schema shifts the burden of inference from encoding to retrieval, requiring the memory system to dynamically reconstruct relationships between events.

Our findings further reveal that memory integration can be supported through multiple pathways. In addition to schema-driven integration, context reinstatement during encoding also contributed to successful inference and was negatively associated with schema reinstatement (**Figure 4A**). This suggests an alternative route to forming unified memory representations supported by event-specific information. In particular, when early schematic reinstatement is weak or absent, later contextual reinstatement may support the integration of overlapping events. Furthermore, context reinstatement predicted subsequent context memory, highlighting its role in supporting detailed event representations (*24*, *37*, *38*). Together, these findings point to a secondary pathway for memory integration, emphasizing high-fidelity context-specific representations rather than abstract schematic knowledge. Speculatively, these two pathways may rely on distinct cognitive and neural mechanisms (*27*, *39*).

Our results further reveal that schema-related processes are dynamically modulated rather than simply engaged or disengaged. During schema-incongruent encoding, suppression of the original incongruent schema was accompanied by the reactivation of the actual schema of Word B, indicating flexible schema processing. Notably, this alternative schema recruitment was associated with reduced reliance on context reinstatement at retrieval (**Figure 4B**), suggesting that when any relevant schema can be recruited during encoding, inference may shift toward integration-based mechanisms. Future work could systematically manipulate competing schemas to determine how alternative knowledge structures are selected and whether initially incongruent schemas are actively suppressed to enable the formation of new integrative representations.

Together, these findings indicate that inference across schema-congruent events is primarily supported by encoding-based integration, but that alternative pathways—such as schema reinstatement at retrieval or context-based integration—can be recruited when encoding-based integration is incomplete, highlighting the adaptive flexibility of memory.

Finally, our findings reveal a dissociation between encoding- and retrieval-related processes in supporting inference. In schema-congruent events, schema reinstatement during encoding played a central role, whereas schema-related activity at retrieval does not simply reflect reactivation of integrated representations, but instead contributes to later stages of inference. In contrast, when events were schema-incongruent inference relied primarily on retrieval-related context reinstatement. Together, these results suggest that encoding and retrieval contribute differently to inference depending on how new experiences relate to prior knowledge.

Although integrative encoding has been associated with faster inference than retrieval-based recombination (*4*, *32*), we did not observe this difference. MVPA results suggest that both conditions included a mixture of representational strategies, which may have obscured differences in response time and masked the expected effect. When analyses were restricted to trials exhibiting the dominant strategy in each condition, the expected response time difference emerged (see **Supplementary Note 4**), consistent with prior work (*4*, *32*).

Previous research has shown that the brain can flexibly form both integrated and separate representations of overlapping experiences to support different task demands (*40*). However, the relative emergence of these representations depends on several boundary conditions, including task demands (*41*), time (*42*), encoding context (*37*, *43*), and individual differences (*32*, *44*). Our findings extend prior work by identifying prior knowledge as a key boundary condition that operates at encoding, biasing whether overlapping experiences are integrated or stored as distinct traces. In doing so, prior knowledge not only influences memory strength, but determines the representational format of memory and the mechanistic route through which inference is later achieved.

Several limitations should be noted. First, although our results are consistent with integrative encoding in schema-congruent conditions, the formation of integrated representations is inferred indirectly rather than measured directly. Second, while EEG provides high temporal resolution for tracking reinstatement dynamics, it does not allow precise localization of the underlying neural sources. Future work combining high temporal and spatial resolution methods will be important for identifying the neural circuits supporting these processes. Finally, our paradigm operationalized prior knowledge in terms of semantic schemas; whether similar mechanisms extend to other forms of prior knowledge remains an open question.

In summary, our findings demonstrate that prior knowledge dynamically shapes memory-based inference. Schema-congruent events are preferentially integrated during encoding, forming unified representations that span multiple experiences. This integrative process likely reflects the embedding of new information within existing schematic frameworks, supported by the medial prefrontal cortex (*33*, *45*). In contrast, schema-incongruent events are more often encoded as distinct memory traces, and the inference across them depends on the flexible retrieval and recombination of these separate traces at test, a process thought to be supported by a big-loop recurrence within the hippocampus (*46*). Together, these findings highlight how prior knowledge shapes inferences and suggest that similar principles may extend to other domains of structured knowledge, including social and cultural schemas that guide decision-making and behavior (*47*).

## METHOD

### Participants

To ensure adequate statistical power, and following previous studies (*32*), we collected data from a total of 40 participants. Data from one participant was excluded due to excessive noise in the EEG. Thus, the final sample consisted of 39 participants (average age 29.9 ± 6.4 years, 10 males and 29 females), all with normal or corrected-to-normal vision. All participants were right-handed, fluent in English, and reported no psychiatric or neurological disorders. For their time in the lab (averaging 3 hours per participant, including EEG preparation), participants received monetary compensation in the form of vouchers to be used in various shops.

All procedures were carried out in accordance with the Swedish Act concerning the Ethical Review of Research involving Humans (2003:460) and the World Medical Association’s Declaration of Helsinki’s Code of Ethics. Participants provided their informed written consent before taking part in the study. The data collection was anonymous and did not involve any potentially identifying personal information. As established by Swedish authorities and specified in the Swedish Act concerning the Ethical Review of Research involving Humans (2003:460), this study does not require specific ethical review by the Swedish Ethical Review Authority for the following reasons: (1) it does not deal with sensitive personal data, (2) it does not use methods that involve physical intervention, (3) it does not use methods that pose a risk of mental or physical harm, (4) it does not study biological material taken from a living or deceased human that can be traced back to that person.

### Stimuli material

A total of 64 color photos, downloaded from various online sources and resized to 1920 × 1080 pixels, served as background contexts. We used eight different contextual pictures for each schema. Half of the schemas were outdoor (city, forest, beach, and farm), and the other half were indoor (kitchen, airport, bathroom, and classroom).

Additionally, 128 images of human faces (half male and half female) taken from the Oslo Face Database (*29*) and 128 images of dogs downloaded from various online sources were used in the experiment to form picture-word associations. All the dog images were removed from their original backgrounds, resized to 500 × 500 pixels, and placed on a light grey surface in a white background to match the face stimuli.

A pool of 128 singular concrete nouns was selected from the Small World of Words dataset for English word associations (*48*). Specifically, we selected 16 congruent words for each schema based on their association with that schema. The frequency of the words was checked against the Corpus of Contemporary American English (*49*). The schema-congruent words were divided into four lists. We tested if the lists were equivalent in terms of association strength with the schema, frequency, and length. This was evaluated with the *BayesianFactor* (0.9.12) implemented in R (4.1.2), the Bayesian Factor in favor of the null hypothesis over the alternative hypothesis (BF_01_) was calculated and a criterion of three was adopted (*50*). We confirmed that the lists were matched in terms of association strength (BF_01_ = 22.611), length (BF_01_ = 5.323), and frequency (BF_01_ = 18.822). Each word was randomly paired with a face and a dog to form the AB and the BC associations. For half of the corresponding AB and BC associations, A was a face, and C was a dog; for the other half, the roles were reversed. Each list was assigned to one experimental condition. The incongruent lists were formed by shuffling words across different schemas, ensuring that the words did not fit the schema (e.g., the word ‘fridge’ and ‘baggage’ for the schema of *beach*). To ensure that differences between conditions were not due to differences in the stimulus material, the assignment of the lists to the experimental conditions was counterbalanced across participants.

### Procedure

The stimuli were presented electronically using the E-Prime 3.0 software (Psychology Software Tools, Inc. [E-Prime Go], 2020) on a 19-in. monitor with a resolution of 1920 ×1080 pixels. The stimuli were presented against a black background. The words appeared in a white rectangular box measuring 100 × 200 pixels, in bold Arial Unicode MS, size 20.

The session started with the localizer task, in which participants were presented with 64 contextual images corresponding to the eight schemas. The pictures were randomly presented, one at a time and for 2000 milliseconds. Participants were asked to identify whether the contextual picture represented an indoor or outdoor scene by pressing ‘i’ for indoor and ‘o’ for outdoor. Each contextual picture was shown five times. The data acquired during this localizer task were used to build the multivariate pattern classifiers of schema and context (see **Figure 1B**).

After the localizer task, participants were introduced to the memory task. The task comprised eight blocks, and participants were encouraged to take breaks between blocks. Each block consisted of an encoding phase, a distractor task—in which participants successively subtracted seven from a random three-digit number for 30 seconds, and a retrieval phase. In the encoding phase, participants were asked to encode events consisting of a word and a picture (face/dog) displayed on a background (see **Figure 1A**). For AB events, the backgrounds are contextual images, which can be either schematically congruent or incongruent with the Word B. In contrast, BC events comprise a black background, with their schema congruency inherited from the corresponding AB events. Participants were asked to learn eight AB events, followed by eight BC events. Non-overlapping XY events were also implemented in the present experiment. Intermixed with AB events, event X was learned, in which a word was paired with a light grey square matching the size of picture A and presented on a contextual background. These were followed by XY events, in which the word X was paired with picture Y (face/dog) and presented on a black background, learned intermixed with BC events. Half of the events were schema-congruent, and the other half were schema-incongruent. In each block, participants encoded congruent and incongruent events of two schemas, each consisting of an indoor and an outdoor environment. The eight schemas were grouped in pairs in the following way: a) beach and classroom, b) city and kitchen, c) forest and airport, and d) farm and bathroom. There were two blocks of each schema pair, and the presentation order of the schema pairs was counterbalanced across participants. Participants were asked to encode all the direct (AB, BC, and XY) and inferred associations (AC) and to memorize the context in which the associations were presented. In the retrieval phase, participants were first tested on all the inferred AC associations, followed by the retrieval of the schema and the context in which picture A appeared. Finally, all direct associations were tested (see **Figure 1A**).

The AB encoding trial started with a fixation cross for 1 second. Next, the contextual picture was displayed for 2 seconds, followed by the Word B presented for 1 second on the right side of the screen. After that, the picture of the face/dog was displayed on the left side of the screen and remained there, along with the word and the context, for another 5 seconds. The BC trials started with the presentation of the C, for 5 seconds, on the left side of the screen, followed by the presentation of the B for another 5 seconds, on the right side of the screen. Each trial was preceded by a jittered black screen, ranging between 0.25 and 0.75 seconds. At retrieval, the trials started by presenting a fixation cross for 1 second. After that, the cue, picture A, was displayed for 2 seconds. Thereafter, the target, corresponding picture C, and a distractor, a picture Y from a non-overlapping XY association, were presented in the bottom half of the screen and remained for a maximum of 6 seconds. The distractor was presented during encoding in the same contextual picture and belongs to the same category (dog vs. face) as the target. The AC test was followed by the schema memory test, in which participants indicated the schema where the A was originally presented (schema memory test). If participants selected the correct schema, they proceeded to the context memory test, indicating the specific context (out of four choices) in which A had been presented. If the schema memory response was incorrect, participants moved directly to the next AC test. The direct association tests comprised the final part of the retrieval phase and followed the same structure as the AC test, except that the cue was the Word B/X.

The accuracy of schema memory is used to control for random guessing in the context memory test. However, this correctness-based control could also provide feedback on schema retrieval, potentially affecting confidence and subsequent direct association tests (i.e., AB and BC). To examine this possibility, we conducted a Bayesian analysis to test whether schema memory performance moderated the effects of congruency and association type. If schema retrieval outcomes influenced performance on the direct association memory tests, this could indicate that the experimental procedure introduced systematic bias. Conversely, if no moderation effect was present, performance on the direct association should be independent of whether participants progressed to the context memory test after the schema memory test. This analysis was performed using the *BayesFactor* package (version 0.9.12). Evidence in favor of the null hypothesis over the alternative hypothesis was quantified using the Bayesian factor (BF_01_), with a threshold of three adopted as the criterion for moderate support for the null over the alternative hypothesis (*50*, *51*). The test result strongly favored the null hypothesis over the alternative (BF_01_ = 1081.794), indicating that the experimental procedure did not bias participants’ performance on direct association tests.

Across all memory tests, participants were given the option to press “I don’t know” if they did not remember, to minimize noise from guessing. “I don’t know” responses were considered as incorrect trials in the analyses. In **Supplementary Note 5**, we summarized, across all participants, the overall proportion of “I don’t know” responses among incorrect trials and examined that their frequency did not vary across experimental conditions.

### EEG data collection and preprocessing

A SynAmps RT Neuroscan amplifier (1 kHz sampling rate; left mastoid reference; bandwidth DC-3500 Hz; 24-bit resolution) with 62 active electrodes mounted in an elastic cap was used to collect EEG data. The electrodes were positioned according to the extended 10-20 system. Additional electrodes were used for references (left and right mastoids), virtual ground (placed between Fpz and Fz), and vertical electrooculogram (EOG, placed in the cheek, below the right eye). The EEG data were preprocessed using FieldTrip (Oostenveld et al., 2011) and in-house MATLAB scripts. Offline, the EEG data was down-sampled to 500 Hz. A lowpass (up to 100 Hz) and a notch filter (50 and 100 Hz) were applied to the data to extract the frequency of interest and remove common noise of alternating current.

Subsequently, the data was segmented into epochs. The data from the localizer task were used to build the hierarchical schema and context classifiers. These data were epoched from −1 to 3 s relative to the onset of the contextual picture. The AB encoding epochs, used to validate the schema and context classifiers, ranged from −1 to 3 seconds relative to the onset of the first context picture without other event elements. The BC encoding epochs ranged from −1 to 5 seconds relative to the onset of the presentation of both B and C. The AC inference test epochs ranged from −1 to 3 seconds relative to the onset of the cue, including the 1-second period when the target and distractor were also presented. The epoched data was transformed into a link-mastoid reference and demeaned (by subtracting the average amplitude of the whole epoch). Additional electrooculogram measures were estimated using the Fp2 electrode and the electrode below the right eye, and the FT7 and FT8 electrodes in the cap to detect vertical (blinks) and horizontal eye movements.

The epochs were visually inspected to remove those with artifacts other than excessive muscle activities, blinks, and eye movements. Independent component analysis was conducted to remove the components representing oculomotor artifacts and muscle activities that were distinct from the EEG signal. In addition to all the above, bad channels were identified before the whole preprocessing and were interpolated before analysis. Finally, the data were again visually inspected to remove trials containing remaining artefacts.

The final analysis included an average of 311 familiarization trials (ranging from 292 to 319 trials), 59 AB trials (ranging from 51 to 64 trials), 61 BC trials (ranging from 57 to 64 trials) per participant. The data from the inference test contained on average 60 AC test trials (ranging from 58 to 63 trials).

### Behavioral data analysis

The behavioral data were analyzed with linear mixed models using R (4.1.2) packages of *lme4* (1.1-34) and *emmeans* (1.8.7) for model fitting and statistical tests and *effectsize* (0.8.9) for effect size estimation. The first level of each model was the trial level, which was clustered in the second level, i.e., the participant level. The significance criterion of *p* < .05 was applied for all statistical analyses. Trials in which participants did not respond within the given time frame or responded “I don’t know” were considered incorrect.

First, behavioral performance for the AC inference test and the direct associations was contrasted in terms of accuracy and response times (RTs). Linear mixed models were fitted with the factors Association Type (AC vs. AB vs. BC vs. XY) and Congruency (Congruent vs. Incongruent). The first level of each model was the trial level, which was clustered in the second level, i.e., the participant level. To rule out potential confounds of random effects (*52*) and simultaneously prevent model overfitting (*53*), the fitting of each model started with participants as the only random intercept. Other factors were thereafter added as random slopes in a step-by-step fashion. If an added random slope significantly improved model fit (i.e., *p* < .05 in a chi-square test for model fit comparison, and a decrease in Akaike Information Criterion and Bayesian Information Criterion), the random slope was retained. The final equations for all models, showing which random factors were considered, are provided in **Supplementary Note 1**. For each model, the homogeneity of variance of the residuals was assessed by using Levene’s test. If the test indicated heteroskedasticity, we re-ran the model with a restricted variance structure (*13*, *37*). For all statistical analyses, the significance criterion of *p* < .05 was applied. Correct responses outside a 10-second time window were considered incorrect to prevent responses based on logical reasoning rather than memory to bias the results. Only correct responses were included in the response-time analysis.

To infer the mechanism underlying memory integration across schema-congruent and incongruent events, we tested the extent to which the AC inference depended on the joint accuracy of AB and BC associations. If AC inference was made by integrating AB and BC memories during encoding, then AC inference should not show reliance on the joint AB and BC accuracy at the test phase. On the other hand, if AC inference is accomplished via flexibly retrieving AB and BC at the time of test, then the joint accuracy of AB and BC should predict AC inference performance. Linear mixed models were constructed using the joint accuracy of AB and BC associations to predict the corresponding AC accuracy, with AB and BC accuracies included as control variables (see **Supplementary Note 1** for the model equation). Separate models were built for schema-congruent and schema-incongruent events.

An additional measure, employing corrected dependency to estimate the integration of AB and BC during encoding (*44*, *54*), was also implemented to assess the integrative encoding of AB and BC events. However, as it did not adequately capture variability in cognitive processes across different schema conditions, the corresponding results are presented in **Supplementary Note 2**.

Schema memory analysis was calculated across all trials; however, for the context memory analysis, only correct schema memory trials were included. Linear mixed models were performed with the factors of Congruency (Congruent vs. Incongruent) and AC accuracy (correct vs. incorrect) to estimate how schema congruency and inference affect memory for schema and context (see **Supplementary Note 1** for model equations).

The normality of the residuals model was assessed by inspecting the Q-Q plots of the standardized residuals and Shapiro-Wilk’s tests. The homogeneity of variance of the model residuals was assessed for each model by visually inspecting a plot of the model residuals versus fitted values and using Levene’s test for unequal variance. Significant complex effects were followed up with post hoc tests with Tukey-correction. Effect sizes, partial eta squared (*η*_*p*_^2^) for F-tests and unstandardized difference (*D*) for t-tests, were also reported together with other statistics.

### Schema and context multivariate pattern classification

The multivariate pattern classification was implemented with the MVPA-light package (*55*) and in-house MATLAB scripts. Multiclass linear discriminant analysis was used to classify the EEG amplitude into different schemas and contexts. The classification at the schema level aimed at classifying the EEG amplitudes into eight different schemas (i.e., beach vs. farm vs. forest vs. city vs. classroom vs. bathroom vs. airport vs. kitchen), while the classification at the context level aimed at classifying the EEG amplitudes into eight different exemplars (contexts) within each schema (see **Figure 1B**). The pattern classifiers were trained on data collected during the localizer task, which included EEG waveforms elicited during viewing each picture five times.

Prior to classification, the EEG waveforms were down-sampled to 100Hz and smoothed with a 0.1s moving window to reduce redundancy in classification features and speed up computations. After that, each epoch was baseline-corrected by subtracting the averaged amplitude between −0.2 and 0s for each channel. Finally, the EEG amplitudes were z-scored across trials. Classifiers were built at each time point of the EEG waveform, between −0.2s to 2s with respect to the onset of the contextual picture and tested on each time point of AB encoding trials to validate the classifiers (see **Figure 1C**), resulting in a time-generalized classification matrix of 221 × 221 time points. The time-generalized classification matrices for all eight context classifiers were averaged to obtain the context classification for each participant.

To investigate the neural substrate of schema and context classification, we examined the contribution of channels to these classifications. We trained classifiers to distinguish schema and context using the localizer data of each channel and their neighboring channels at each time point (*31*), which was tested with the same channels at the same time point of the AB data. This procedure produced a time course of accuracy estimates for each channel, showing how each channel contributes to classifying schema and context over time. The topography of the channel contribution is summarized in **Figure 1D**.

Next, the schema and context classifiers were tested on all time points during BC encoding and during the AC test to investigate schema and context reinstatement, resulting in a time-generalization matrix of 221 × 521 for BC encoding data and 221 × 221 for AC inference test data). The time-generalized classification matrices for all eight context classifiers were averaged to obtain the context classification accuracy for each participant.

During BC encoding, Word B could theoretically elicit schema-like neural responses, potentially contaminating the results—particularly the schema reinstatement measure. A control analysis was therefore conducted (see **Supplementary Note 3)**. The results confirmed that for BC encoding trials, the schema classifiers captured neural activity predominantly associated with the previously encoded AB event, reflecting inference-related demands rather than simple responses to the on-screen word.

### Statistical analysis of the multivariate pattern classification

To reduce the impact of multiple comparison issue in classic statistics, the statistical significance of the classification accuracy was tested with Bayesian statistics using a Beta distribution prior (alpha = 30, beta = 210) that accounts for potential false discoveries by applying constraints in prior expectation and precision (*32*, *56*). For the validation with AB encoding trials, the overall results were obtained by accumulating the accuracy of each participant with consecutive Bayesian update, and the posterior was contrasted against chance (1/8). For the schema and context reinstatement capture in BC encoding and AC test trials, the classification accuracies were contrasted between trials with correct AC inference versus trials with incorrect AC inference (*57*), to identify schema and context reinstatement that contributed to AC inference. To further examine our interpretations, we also tested schema and context reinstatement separately for AC-correct and AC-incorrect trials. In this case, the obtained classification was contrasted against chance (1/8). The overall estimate across participants was obtained by sequentially updating the Bayesian posterior with each participant’s difference scores. The final posterior distribution was compared against 0. For all time-generalized analyses, the Bayesian Factor in favor of the alternative hypothesis (BF_10_) > 3 (*50*) was adopted as the indicator of significance. Only significant results lasting for at least 150 ms are reported to avoid spurious findings. For visualization purposes, classification result plots were smoothed using a 2D Gaussian kernel (50 ms width). Importantly, smoothing was not used when evaluating the statistical significance of the effects or when reporting the results.

Given our predictions that memory integration for schema-congruent and incongruent events is supported by different mechanisms, the statistical tests of the EEG data were performed separately for schema-congruent and incongruent trials.

### Relationship between schema and context reinstatement and memory

After identifying time windows of schema and context reinstatement during the BC encoding and AC test, we further examined whether they represent distinct processes or reflect different aspects of the same process by evaluating their co-occurrence in trials with correct AC inference. If they tend to co-occur in the same trials, i.e., show significant positive correlation on a trial basis, they are more likely to reflect different aspects of the same process; if they tend to be observed in different trials, i.e., show significant negative correlation on a trial basis, then they might indicate different, complementary processes that jointly lead to successful AC inference. A Bayesian linear regression with an Inverse-Gamma-Normal distribution prior (alpha = 15, beta = 15, mu = 0, lambda = 30 × *I*, where *I* denotes the identity matrix) was applied to the trial-level data, accounting for false discovery rate correction. Results for all participants were obtained through consecutive Bayesian updates. The standardized regression coefficient was evaluated to reduce the influences of individual differences, the posterior of which was contrast against 0, and the BF_10_ > 3 was also adopted as the indication of significance.

Furthermore, to assess the potential effect of schema and context reinstatement on memory for schema and context, we used these reinstatement measures to predict participants’ performance in the memory tests. This was done using Bayesian linear regression on data at the trial level. The prior, procedure, and statistical criteria remained the same as above.

## Data and Code Accessibility

Data and code will be made available on request.

## Author Contribution

ZL, MJ & IB – Conceptualization and Methodology; ZL – Investigation, Data Curation and Formal Analysis; ZL & IB – Visualization, Writing – Original Draft Preparation; ZL, MJ & IB – Writing – Review & Editing; IB & MJ – Supervision & Funding Acquisition.

## Acknowledgements

This work was supported by the Swedish Research Council Grant VR 2019-02455. We thank Ieva Valavičiūtė for the assistance in data collection and stimuli preparation and all the volunteers who participated in this study. The computations were enabled by resources provided by LUNARC, The Centre for Scientific and Technical Computing at Lund University.

## REFERENCES

1. M. L. Schlichting, A. R. Preston, Memory integration: neural mechanisms and implications for behavior. Curr. Opin. Behav. Sci. 1, 1–8 (2015).

2. M. T. R. Van Kesteren, D. J. Ruiter, G. Fernández, R. N. Henson, How schema and novelty augment memory formation. Trends Neurosci. 35, 211–219 (2012).

3. H. Eichenbaum, The hippocampus and mechanisms of declarative memory. Behav. Brain Res. 103, 123–133 (1999).

4. D. Shohamy, A. D. Wagner, Integrating memories in the human brain: hippocampal-midbrain encoding of overlapping events. Neuron 60, 378–389 (2008).

5. D. Zeithamova, M. L. Schlichting, A. R. Preston, The hippocampus and inferential reasoning: building memories to navigate future decisions. Front. Hum. Neurosci. 6, 1–14 (2012).

6. D. Kumaran, D. Hassabis, J. L. McClelland, What Learning Systems do Intelligent Agents Need? Complementary Learning Systems Theory Updated. Trends Cogn. Sci. 20, 512–534 (2016).

7. D. Kumaran, J. L. McClelland, Generalization through the recurrent interaction of episodic memories: A model of the hippocampal system. Psychol. Rev. 119, 573–616 (2012).

8. J. L. McClelland, B. L. McNaughton, R. C. O’Reilly, Why There Are Complementary Learning Systems in the Hippocampus and Neocortex:InsightsFrom the Successesand Failuresof Connectionist Models of Learning and Memory. 102, 419–457 (1995).

9. R. C. O’Reilly, J. W. Rudy, Computational principles of learning in the neocortex and hippocampus. Hippocampus 10, 389–397 (2000).

10. V. E. Ghosh, A. Gilboa, What is a memory schema? A historical perspective on current neuroscience literature. Neuropsychologia 53, 104–114 (2014).

11. G. Brod, U. Lindenberger, M. Werkle-Bergner, Y. L. Shing, Differences in the neural signature of remembering schema-congruent and schema-incongruent events. NeuroImage 117, 358–366 (2015).

12. J. R. Anderson, Effects of prior knowledge on memory for new information. Mem. Cognit. 9, 237–246 (1981).

13. S. Audrain, M. P. McAndrews, Schemas provide a scaffold for neocortical integration of new memories over time. Nat. Commun. 13, 5795 (2022).

14. B. P. Staresina, J. C. Gray, L. Davachi, Event Congruency Enhances Episodic Memory Encoding through Semantic Elaboration and Relational Binding. Cereb. Cortex 19, 1198–1207 (2009).

15. M. T. R. Van Kesteren, P. Rignanese, P. G. Gianferrara, L. Krabbendam, M. Meeter, Congruency and reactivation aid memory integration through reinstatement of prior knowledge. Sci. Rep. 10, 4776 (2020).

16. A. Greve, E. Cooper, A. Kaula, M. C. Anderson, R. Henson, Does prediction error drive one-shot declarative learning? J. Mem. Lang. 94, 149–165 (2017).

17. M. T. R. Van Kesteren, S. F. Beul, A. Takashima, R. N. Henson, D. J. Ruiter, G. Fernández, Differential roles for medial prefrontal and medial temporal cortices in schema-dependent encoding: From congruent to incongruent. Neuropsychologia 51, 2352–2359 (2013).

18. D. Frank, D. Montaldi, B. Wittmann, D. Talmi, Beneficial and detrimental effects of schema incongruence on memory for contextual events. Learn. Mem. 25, 352–360 (2018).

19. J. Ortiz-Tudela, G. Turan, M. Vilas, L. Melloni, Y. L. Shing, Schema-driven prediction effects on episodic memory across the lifespan. Philos. Trans. R. Soc. B Biol. Sci. 379, 20230401 (2024).

20. E. Tulving, N. Kroll, Novelty assessment in the brain and long-term memory encoding. Psychon. Bull. Rev. 2, 387–390 (1995).

21. R. A. Diana, A. P. Yonelinas, C. Ranganath, Parahippocampal cortex activation during context reinstatement predicts item recollection. J. Exp. Psychol. Gen. 142, 1287–1297 (2013).

22. J. F. Miller, M. Neufang, A. Solway, A. Brandt, M. Trippel, I. Mader, S. Hefft, M. Merkow, S. M. Polyn, J. Jacobs, M. J. Kahana, A. Schulze-Bonhage, Neural Activity in Human Hippocampal Formation Reveals the Spatial Context of Retrieved Memories. Science 342, 1111–1114 (2013).

23. S. Folkerts, U. Rutishauser, M. W. Howard, Human Episodic Memory Retrieval Is Accompanied by a Neural Contiguity Effect. J. Neurosci. 38, 4200–4211 (2018).

24. I. Bramão, J. Jiang, A. D. Wagner, M. Johansson, Encoding contexts are incidentally reinstated during competitive retrieval and track the temporal dynamics of memory interference. Cereb. Cortex 32, 5020–5035 (2022).

25. S. DuBrow, N. Rouhani, Y. Niv, K. A. Norman, Does mental context drift or shift? Curr. Opin. Behav. Sci. 17, 141–146 (2017).

26. C. Baldassano, U. Hasson, K. A. Norman, Representation of Real-World Event Schemas during Narrative Perception. J. Neurosci. 38, 9689–9699 (2018).

27. R. Masís-Obando, K. A. Norman, C. Baldassano, Schema representations in distinct brain networks support narrative memory during encoding and retrieval. eLife 11, e70445 (2022).

28. S. H. P. Collin, R. P. Kempner, S. Srivatsan, K. A. Norman, Neural codes track prior events in a narrative and predict subsequent memory for details. Commun. Psychol. 3, 26 (2025).

29. O. Chelnokova, B. Laeng, M. Eikemo, J. Riegels, G. Løseth, H. Maurud, F. Willoch, S. Leknes, Rewards of beauty: the opioid system mediates social motivation in humans. Mol. Psychiatry 19, 746–747 (2014).

30. J.-R. King, S. Dehaene, Characterizing the dynamics of mental representations: the temporal generalization method. Trends Cogn. Sci. 18, 203–210 (2014).

31. I. Bramão, M. Johansson, Neural Pattern Classification Tracks Transfer-Appropriate Processing in Episodic Memory. eneuro 5, ENEURO.0251-18.2018 (2018).

32. I. Bramão, Z. Liu, M. Johansson, Remembering the past affects new learning: The temporal dynamics of integrative encoding. Neuropsychologia 212, 109148 (2025).

33. A. Gilboa, H. Marlatte, Neurobiology of Schemas and Schema-Mediated Memory. Trends Cogn. Sci. 21, 618–631 (2017).

34. M. L. Schlichting, A. R. Preston, Memory reactivation during rest supports upcoming learning of related content. Proc. Natl. Acad. Sci. 111, 15845–15850 (2014).

35. D. Zeithamova, A. R. Preston, Flexible memories: differential roles for medial temporal lobe and prefrontal cortex in cross-episode binding. J. Neurosci. 30, 14676–14684 (2010).

36. A. Tompary, W. Zhou, L. Davachi, Schematic memories develop quickly, but are not expressed unless necessary. Sci. Rep. 10, 16968 (2020).

37. Z. Liu, M. Johansson, R. Johansson, I. Bramão, The effects of episodic context on memory integration. Sci. Rep. 14, 30159 (2024).

38. C. Ranganath, Binding Items and Contexts: The Cognitive Neuroscience of Episodic Memory. Curr. Dir. Psychol. Sci. 19, 131–137 (2010).

39. L. A. Libby, Z. M. Reagh, N. R. Bouffard, J. D. Ragland, C. Ranganath, The hippocampus generalizes across memories that share item and context information. J. Cogn. Neurosci. 31, 24–35 (2019).

40. Z. Liu, M. Johansson, I. Bramão, Episodic events are flexibly encoded in both integrated and separated neural representations. Nat. Commun. 17, 752 (2026).

41. F. R. Richter, A. J. H. Chanales, B. A. Kuhl, Predicting the integration of overlapping memories by decoding mnemonic processing states during learning. NeuroImage 124, 323–335 (2016).

42. D. Zeithamova, A. R. Preston, Temporal proximity promotes integration of overlapping events. J. Cogn. Neurosci. 29, 1311–1323 (2017).

43. W. R. Cox, S. Dobbelaar, M. Meeter, M. Kindt, V. A. Van Ast, Episodic memory enhancement versus impairment is determined by contextual similarity across events. Proc. Natl. Acad. Sci. 118, e2101509118 (2021).

44. W. Yu, K. D. Duncan, M. L. Schlichting, Using retrieval contingencies to understand memory integration and inference. Mem. Cognit., doi: 10.3758/s13421-025-01727-8 (2025).

45. O. Bein, Y. Niv, Schemas, reinforcement learning and the medial prefrontal cortex. Nat. Rev. Neurosci. 26, 141–157 (2025).

46. R. Koster, M. J. Chadwick, Y. Chen, D. Berron, A. Banino, E. Düzel, D. Hassabis, D. Kumaran, Big-Loop Recurrence within the Hippocampal System Supports Integration of Information across Episodes. Neuron 99, 1342–1354.e6 (2018).

47. M. Boeltzig, M. Johansson, I. Bramão, Ingroup sources enhance associative inference. Commun. Psychol. 1, 40 (2023).

48. S. De Deyne, D. J. Navarro, A. Perfors, M. Brysbaert, G. Storms, The “Small World of Words” English word association norms for over 12,000 cue words. Behav. Res. Methods 51, 987–1006 (2019).

49. M. Davies, Corpus of Contemporary American English, Corpus of Contemporary American English (2008). https://www.english-corpora.org/coca/.

50. R. E. Kass, A. E. Raftery, Bayes factors. J. Am. Stat. Assoc. 90, 773–795 (1995).

51. F. D. Schönbrodt, E.-J. Wagenmakers, Bayes factor design analysis: Planning for compelling evidence. Psychon. Bull. Rev. 25, 128–142 (2018).

52. W. T. Ambrosius, Ed., Topics in Biostatistics (Humana Press, Totowa, N.J, 2007)Methods in molecular biology.

53. H. Matuschek, R. Kliegl, S. Vasishth, H. Baayen, D. Bates, Balancing Type I error and power in linear mixed models. J. Mem. Lang. 94, 305–315 (2017).

54. A. J. Horner, N. Burgess, The associative structure of memory for multi-element events. J. Exp. Psychol. Gen. 142, 1370–1383 (2013).

55. M. S. Treder, MVPA-Light: A Classification and Regression Toolbox for Multi-Dimensional Data. Front. Neurosci. 14, 289 (2020).

56. H. Han, Implementation of Bayesian multiple comparison correction in the second-level analysis of fMRI data: With pilot analyses of simulation and real fMRI datasets based on voxelwise inference. Cogn. Neurosci. 11, 157–169 (2020).

57. T. Pham-Gia, N. Turkkan, P. Eng, Bayesian analysis of the difference of two proportions. Commun. Stat. - Theory Methods 22, 1755–1771 (1993).

