## Supplementary Note for "Prior knowledge shapes neural routes to novel inference across events"

### Supplementary Note 1: Models for Assessing Behavioral Performance

To assess participants' task performance, we employed linear mixed-effects models to evaluate the impact of variables of interest at the trial level, with individual differences accounted for by random slopes. For all statistical models, to prevent model overfitting (*1*, *2*), each model started with the least complexity, i.e., with participants as the single random intercept. The inclusion of other effects as random slopes was evaluated by examining model fit. If the model fitting improved with the inclusion of a given random slope, the random slope was kept. The final equations for each model are displayed in **Table S1**.

**Table S1** Models for behavioral performance assessment.

| | Equation |
|---|---|
| Associative Memory | |
| | $accuracy \sim Association\ Type + Congruency + Association\ Type * Congruency + intercept + (intercept \mid participant)$ |
| | $response\ times \sim Association\ Type + Congruency + Association\ Type * Congruency + intercept + (intercept \mid participant)$ |
| Schema Memory | |
| | $accuracy \sim AC\ inference + Congruency + AC\ inference * Congruency + intercept + (intercept \mid participant)$ |
| Context Memory | |
| | $accuracy \sim AC\ inference + Congruency + AC\ inference * Congruency + intercept + (intercept \mid participant)$ |
| Dependency of AC Inference on Simultaneous AB and BC Retrieval | |
| | $AC\ inference \sim AB\ accuracy + BC\ accuracy + AB\ accuracy * BC\ accuracy + intercept + (intercept \mid participant)$ |

**Supplementary Note 2: Schema Classifiers Selectively Track Schema Reinstatement Rather Than Word-Driven Neural Activity**

In the present study, schema classifiers trained during the localizer task were used to detect schema reinstatement during BC encoding and AC retrieval, enabling us to assess its contribution to AC test performance. However, the semantic meanings of Word B/X could, in principle, also evoke schema-like neural representations, potentially confounding this measure. To address this possibility, we conducted additional control analyses. Notably, all trials involved were associated with correct XY associations (for XY trials) or AC associations (for BC trials), which are expected to best reflect the underlying cognitive processes.

The analyses comprise two stages. In the first stage, we assessed neural decoding of schema representations for X events (a control condition learned alongside AB events, in which only a word is encoded within a context) during XY trials. If schema classifiers track the semantic content of the on-screen word X, schema-congruent XY trials should show above-chance decoding, whereas schema-incongruent XY trials should show below-chance decoding, reflecting a mismatch between the word X and the schema. In contrast, if schema classifiers track X event schemas reactivated by reencountering word X during XY trials, both trial types should yield above-chance decoding of the X event schema.

The schema classification patterns for schema-congruent and incongruent XY trials were evaluated separately, and were contrasted against chance level (12.5%) using Bayesian statistics (Gamma prior: alpha = 30, beta = 210) (*3*, *4*). Both trial types showed above-chance decoding of the X event schema (**Figure S1B**), suggesting that schema classifiers track reinstatement of X events rather than the semantic content of the on-screen word.

Next, we contrasted the reinstatement of AB and X event schemas in BC and XY trials (**Figure S1C**) to provide complementary evidence of how the brain leverages schema representations to support novel inference across episodic events. We used Bayesian tests to compare BC versus XY trials under two schema conditions (Gamma prior for both, alpha = 30, beta = 210). Schema-congruent BC encoding showed more positive classification compared with schema-congruent XY encoding (accuracy difference = 0.025, $BF_{10}$s > 3.005), indicating stronger reinstatement of AB event schema compared with X event schema, a pattern reflecting inference-related processing. In contrast, schema-incongruent BC encoding exhibited greater negative classification than XY encoding (accuracy difference = -0.024, $BF_{10}$s > 3.005), which implies that AB-schema deactivation is driven by inference demands rather than by simple word-schema incongruency. Together, these results indicate that the effects observed in this study reflect the requirements of inference making rather than differences in semantic processing of the on-screen word.

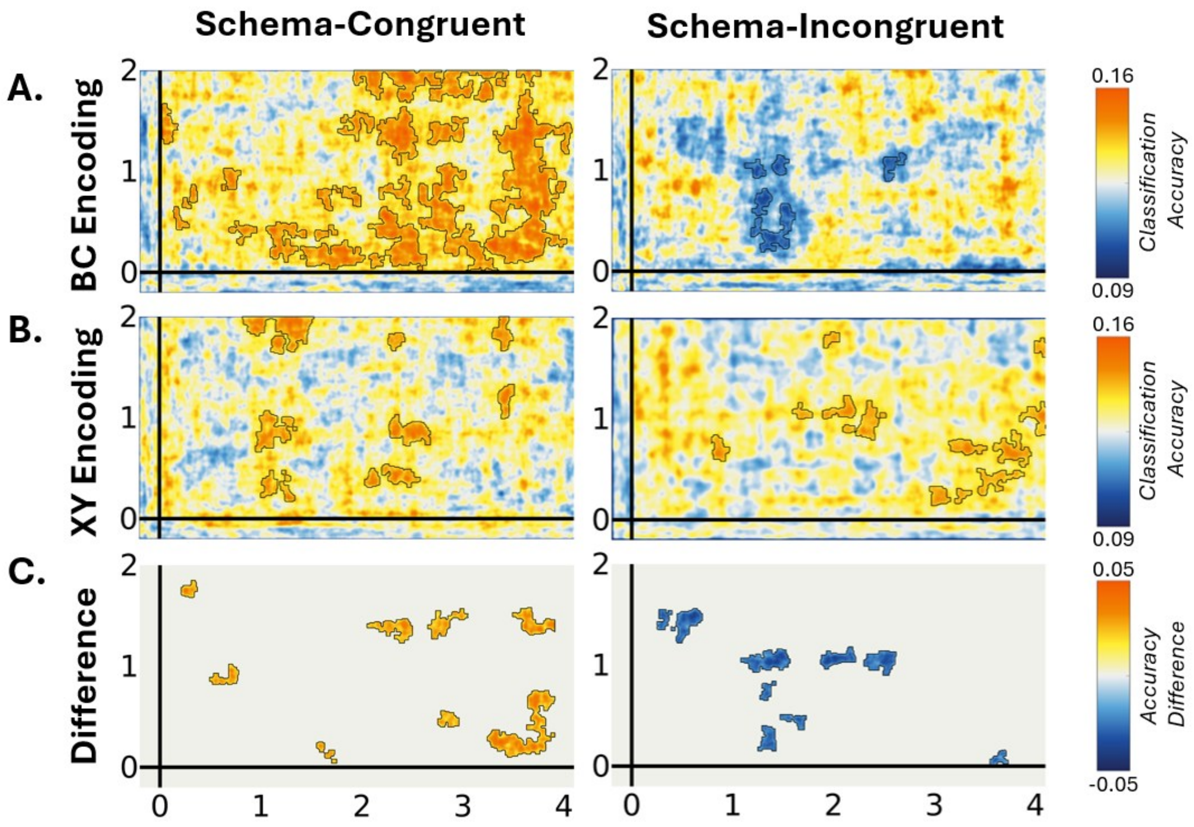


**Figure S1** Time-generalized matrices for schema classification during BC and XY learning, including only trials associated with correct AC or XY associations. For BC and XY encoding, regions with classification accuracy significantly different from chance are indicated by black contours. Direct comparisons between BC and XY encoding for both schema conditions are also shown; areas highlighted exhibit significant differences.

In summary, our analysis demonstrated that the schema classifiers effectively captured genuine schema representations while remaining robust to potential artifacts arising from the semantic meaning of Word B/X. Moreover, the findings indicate that the reinstatement and deactivation of encoded schemas result from the task demands of inference making. Specifically, in the XY conditions—where no cross-episode inference is required—we observed lower schema reinstatement in the schema-congruent condition and no schema deactivation in the schema-incongruent condition. These results further show that the schema classifiers successfully distinguished between encoding a novel event that overlaps with a prior one and encoding a baseline event, underscoring both the efficacy of our methodology and the flexibility of human cognition.

### Supplementary Note 3: Reexamination of Response Times for AC Indirect Associations across Schema-Congruent and Schema-Incongruent Events

Previous studies have shown that response times in AC tests are faster following integrative encoding than flexible retrieval, as the former involves retrieval of an already integrated representation, whereas the latter requires retrieval and recombination of separate memory traces (*5*–*7*).

Neural evidence from our study showed that schema-congruent events are primarily integrated during encoding, forming unified representations that span multiple experiences, whereas schema-incongruent events are more often encoded as distinct memory traces, and the inference across them depends on the flexible retrieval and recombination of separated memory traces on demand. Interestingly, our behavioral analysis revealed only a numerical difference in response times for indirect AC associations between schema-congruent and schema-incongruent conditions, which does not reach statistical significance.

Notably, although relatively uncommon, some schema-congruent events are flexibly recombined during retrieval, while some schema-incongruent events are integrated during encoding (**Figures 3-6**). This unexpected mixture of processes may have blurred the distinction between the two conditions, providing a compelling explanation for the current results and motivating the focused analysis presented here. To test this, we selected a subset of trials from these two conditions that exhibited typical patterns of integrative encoding or flexible retrieval and reexamined the response time differences.

Specifically, for each participant, in the schema-congruent condition, trials were sorted based on the difference between average schema and context reinstatement during BC encoding and the schema reinstatement during AC test (see **Figure 3** for the patterns), and those above the median were selected, reflecting higher engagement of integrative encoding. Conversely, for the schema-incongruent condition, trials were sorted based on the difference between average context reinstatement during the AC test and the reinstatement of the Word B congruent schema during BC encoding (see **Figure 5** and **6** for the patterns), and those above the median were included, indicating greater engagement of flexible retrieval. We then compared response times for correct AC inferences between these two conditions using Bayesian linear regression (Normal-Inverse-Gamma prior, alpha = 15, beta = 15, mu = 0, lambda = 30**I*, *I* is unit matrix), with the reinstatement strength (indicated by average classification accuracy), as well as their interaction term, as control variables. Results showed that, for trials exhibiting typical neural activity patterns of integrative encoding or flexible retrieval, AC inference was significantly faster for schema-congruent than schema-incongruent conditions ($\beta$ = -.155, $BF_{10}$ = 3.935). Additionally, as expected, a higher reinstatement level was associated with faster response time ($\beta$ = -.133, $BF_{10}$ = 10.550), whereas the interaction term was not significant ($\beta$ = -.053, $BF_{10}$ = 1.320).

This finding is consistent with previous research suggesting that retrieval of already integrated representations requires less time than the flexible retrieval and recombination of separately encoded memory traces (*5–7*). Moreover, higher levels of reinstatement were also predictive of faster response times, aligning with prior work demonstrating that reinstatement level, as captured by neural pattern classifiers, serves as a reliable indicator of memory trace strength (*8*, *9*). These results further validate our methodological pipeline in tracing mnemonic processes that support complex cognitive functions. Furthermore, this analysis indicates that response time may not always reliably differentiate between integrative encoding and flexible retrieval at the participant level, as these processes could fluctuate across individual trials, potentially influenced by the experimental design or other modulating factors.

**Supplementary Note 4: Examination of the Proportion of 'I Don't Know' Responses**

During the test phase, participants were encouraged to indicate 'I don't know' when unsure of the correct response. This approach was intended to minimize noise from guessing responses. Given that (a) the proportion of 'I don't know' responses was substantial (see **Table S1**), and (b) the current study does not focus on participants' metacognitive awareness, these responses were treated as incorrect trials in all analyses.

We examined whether the proportion of 'I don't know' responses among the incorrect trials varied based on congruency (congruent vs. incongruent) and association type (AC vs. AB vs. BC vs. XY). For the schema and context memory tests, congruency was the only factor entered into the model. There was no significant main effect of congruency or association type, nor any interaction effects on the proportion of 'I don't know' responses ($p$s > .110). Similarly, congruency did not affect the proportion of 'I don't know' responses in the schema and context memory tests ($p$s > .135). These results indicate that "I don't know" responses did not systematically vary across conditions and therefore did not bias task performance.

**Table S2** Proportions of 'I don't know' responses among incorrect trials across all participants ($M \pm SE$).

| | Congruent | Incongruent |
|---|---|---|
| AC | 0.305 ± 0.046 | 0.317 ± 0.046 |
| AB | 0.341 ± 0.050 | 0.376 ± 0.049 |
| BC | 0.338 ± 0.048 | 0.376 ± 0.048 |
| XY | 0.352 ± 0.048 | 0.358 ± 0.047 |
| Schema | 0.337 ± 0.048 | 0.377 ± 0.048 |
| Context | 0.111 ± 0.023 | 0.084 ± 0.023 |

### REFERENCES


1. H. Matuschek, R. Kliegl, S. Vasishth, H. Baayen, D. Bates, Balancing Type I error and power in linear mixed models. *J. Mem. Lang.* **94**, 305–315 (2017).

2. Z. Liu, M. Johansson, R. Johansson, I. Bramão, The effects of episodic context on memory integration. *Sci. Rep.* **14**, 30159 (2024).

3. E.-J. Wagenmakers, T. Lodewyckx, H. Kuriyal, R. Grasman, Bayesian hypothesis testing for psychologists: A tutorial on the Savage–Dickey method. *Cognit. Psychol.* **60**, 158–189 (2010).

4. T. Pham-Gia, N. Turkkan, P. Eng, Bayesian analysis of the difference of two proportions. *Commun. Stat. - Theory Methods* **22**, 1755–1771 (1993).

5. I. Bramão, Z. Liu, M. Johansson, Remembering the past affects new learning: The temporal dynamics of integrative encoding. *Neuropsychologia* **212**, 109148 (2025).

6. M. L. Schlichting, D. Zeithamova, A. R. Preston, CA1 subfield contributions to memory integration and inference. *Hippocampus* **24**, 1248–1260 (2014).

7. D. Shohamy, A. D. Wagner, Integrating memories in the human brain: hippocampal-midbrain encoding of overlapping events. *Neuron* **60**, 378–389 (2008).

8. J. F. Danker, A. Tompary, L. Davachi, Trial-by-Trial Hippocampal Encoding Activation Predicts the Fidelity of Cortical Reinstatement During Subsequent Retrieval. *Cereb. Cortex*, bhw146 (2016).

9. J. D. Johnson, M. D. Rugg, Recollection and the Reinstatement of Encoding-Related Cortical Activity. *Cereb. Cortex* **17**, 2507–2515 (2007).
